# Lack of evidence for gene-level convergence linked to evolutionary shifts in torpor among placental mammals

**DOI:** 10.1101/2025.11.18.689136

**Authors:** Dimitrios - Georgios Kontopoulos, Alexis-Walid Ahmed, Bernhard Bein, Danielle L. Levesque, Michael Hiller

**Affiliations:** Senckenberg Research Institute, Frankfurt am Main, Germany; Department of Ecology and Evolutionary Biology, University of California, Los Angeles, CA, USA; Institute of Cell Biology and Neuroscience, Faculty of Biosciences, Goethe University Frankfurt, Frankfurt am Main, Germany; School of Biology and Ecology, University of Maine, Orono, ME, USA

**Keywords:** torpor, hibernation, placental mammals, comparative genomics, convergent evolution

## Abstract

Torpor is a key survival strategy that many avian and mammalian lineages evolved in response to challenging environmental conditions. Whether the independent evolution of torpor in different lineages involved changes in the same genes remains poorly understood. Here, we performed comparative screens across 190 placental mammal genomes to comprehensively examine associations between loss, positive selection and evolutionary rate shifts in individual protein-coding genes and evolutionary shifts in torpor use. We find that gene-torpor associations are highly clade-specific, with no gene being able to explain the majority of torpor shifts across the phylogeny of placental mammals. Instead, a relatively higher but limited extent of evolutionary convergence can be detected at the pathway level. Our results suggest that torpor emerged through several genetic routes in placental mammals, which likely explains the vast diversity of torpor use patterns that can be observed among torpor-capable species today.

**Significance:** A fundamental question in evolutionary genomics is whether independent gains and losses of a convergent trait may be achieved through similar or distinct genetic paths. Here, we address this question by focusing on the evolution of torpor across placental mammals, a key trait that likely emerged independently in several lineages. By conducting multiple comparative genomics screens across 190 placental mammal species, we find that individual protein-coding genes have a low explanatory power for evolutionary shifts in torpor. In contrast, there is slightly stronger evidence for pathway-level convergence. Our findings suggest that the wide diversity of patterns of torpor use across placental mammals may, in part, be due to the existence of multiple genetic paths to torpor.

## Introduction

Extant mammals and birds are whole-body endotherms, that is, they can maintain a high and relatively stable body temperature through endogenous production of metabolic heat. Whole-body endothermy comes with several advantages (e.g., biochemical reaction rates that are largely independent of environmental temperature), but is very energetically expensive (Clarke and Pörtner 2010; Grigg et al. 2022). The high energy demands of maintaining a constant body temperature over a range of ambient temperatures may be impossible to meet under certain circumstances, such as during periods of limited availability of food resources. To survive in challenging environments, several mammal and bird species (including bats, rodents, hummingbirds, and doves) can greatly decrease their metabolic rate (and, through this, their body temperature) in a controlled and fully reversible manner—a process known as torpor (Ruf and Geiser 2015; Nowack et al. 2020). Although the characteristics of torpor expression in individual species (e.g., depth, length, frequency, external triggers) vary remarkably across endotherms (Ruf and Geiser 2015; Nowack et al. 2020; Kontopoulos et al. 2025), torpor has been classically subdivided into two broad subtypes: daily torpor (lasting less than 24 hours) and prolonged torpor or hibernation (lasting up to an entire year; Ruf and Geiser 2015; Nowack et al. 2020).

We recently showed that torpor patterns in endotherms generally lie along an evolutionary continuum, where macroevolutionary shifts between no torpor and hibernation typically involve an intermediate step, the use of daily torpor (Kontopoulos et al. 2025). This intermediate step may be transient over evolutionary timescales (e.g., lasting only a few million years) and, hence, occasionally unobserved when reconstructing ancestral torpor states based on information from a small number of extant species. Moreover, the ability to undergo torpor appears to be a convergently-evolved trait that has been gained—and secondarily lost—several times in different endothermic lineages. This includes placental mammals, whose last common ancestor is unlikely to have used torpor (Kontopoulos et al. 2025).

The frequently occurring shifts along the evolutionary torpor continuum prompt the question of whether the evolution of torpor in independent lineages involved changes in the same genomic elements or whether torpor evolved through multiple genetic routes. In support of the former, a few studies uncovered patterns of convergent evolution in the same regulatory or protein-coding regions among mammalian hibernators from different clades (Ferris and Gregg 2019; Nakayama and Makino 2024; Drabeck et al. 2025; Ferris et al. 2025; Steinwand et al. 2025). The number of hibernators included in such studies tends to be relatively low (between four and fourteen), but more worryingly, such studies group together torpor-incapable species with species that use daily torpor (daily heterotherms). The latter practice is potentially problematic from the standpoint of an evolutionary continuum in torpor (Kontopoulos et al. 2025), which implies that evolutionary thresholds (that are unobserved but estimable) separate torpor-incapable species from daily heterotherms and daily heterotherms from hibernators. As we previously showed, the evolutionary boundaries among the three torpor groups (lack of torpor, daily torpor, and hibernation) are weak, whereas species that belong to the same group can greatly differ in their position along the evolutionary continuum (Kontopoulos et al. 2025). For example, some daily heterotherms are phenotypically closer (with respect to the evolutionary torpor continuum) to torpor-incapable species, whereas other daily heterotherms are phenotypically closer to hibernators (Kontopoulos et al. 2025). Thus, the extent of evolutionary convergence at the molecular level throughout the evolutionary torpor continuum has yet to be thoroughly and appropriately explored.

In the present study, we thoroughly examine the associations between individual protein-coding genes and evolutionary shifts in torpor across 190 species of placental mammals. More specifically, we combine classifications of torpor use from the literature with comprehensive comparative genomics screens for signatures of gene loss, positive selection, and changes in the rate of molecular evolution. These enable us to shed new light on the generality of the molecular underpinnings of torpor.

## Results and Discussion

### Data sources

For the present study, we extracted classifications of the longest reported torpor duration (no torpor / daily torpor / prolonged torpor or hibernation) for 190 species of placental mammals for which sequenced genomes were available (Table S1) from the 1,338-species dataset of Kontopoulos et al. (2025). We also extracted reliable ancestral state estimates of torpor (posterior probability > 0.95 according to Kontopoulos et al. (2025)) for internal nodes of the phylogeny (Fig. 1). Our dataset comprises 128 torpor-incapable extant species, 28 daily heterotherms, and 34 hibernators. We then coded the torpor patterns of placental mammals in three different ways:

**i) torpor continuum** (three ordered states of no torpor, daily torpor, and hibernation).
**ii) binary torpor** (no torpor *versus* either daily torpor or hibernation).
**iii) hibernation** (no torpor or daily torpor *versus* hibernation).

**Fig. 1.**
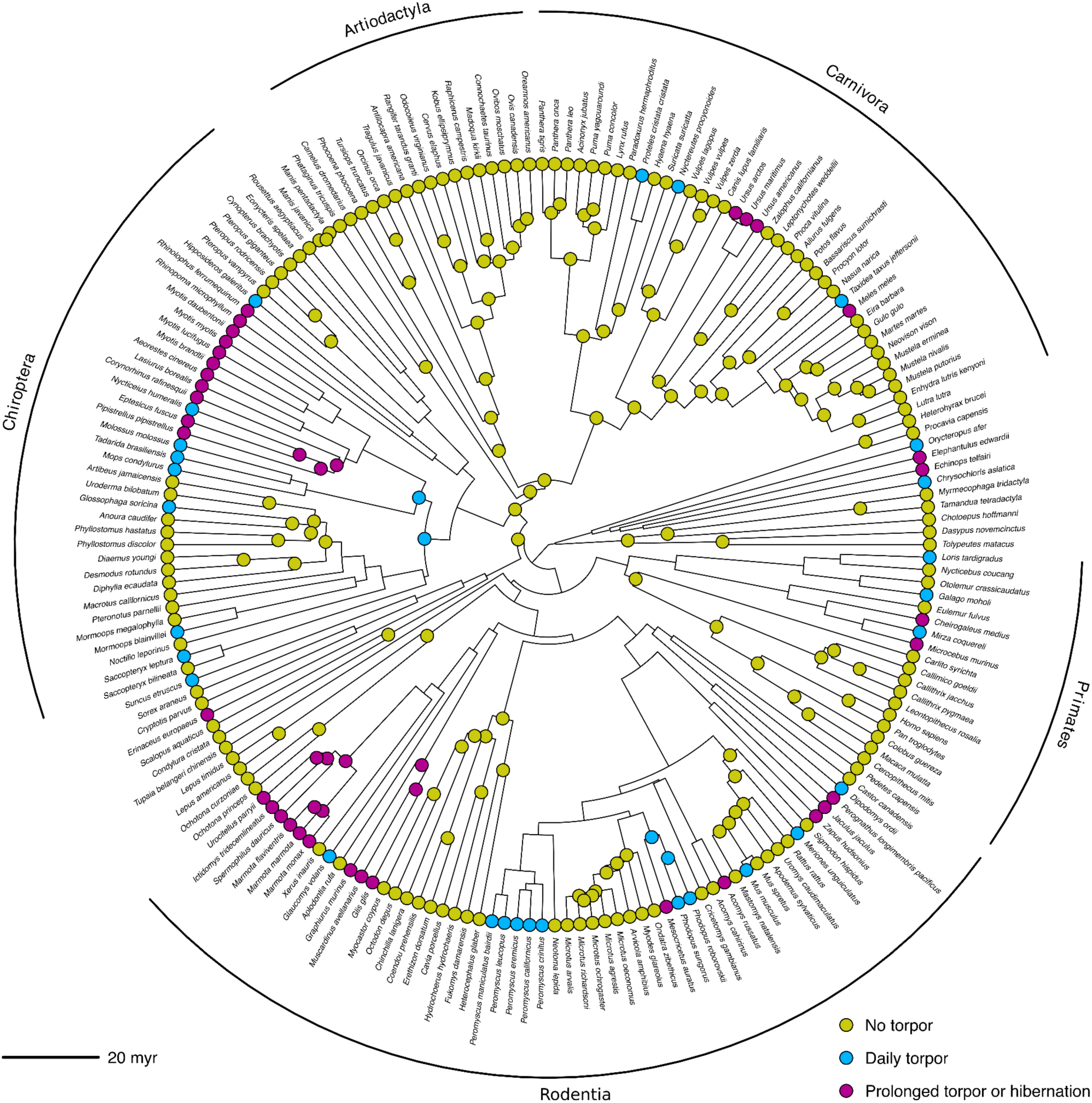
Distribution of torpor use across the 190 species of placental mammals included in our study. The most species-rich orders in our dataset are explicitly shown. All torpor classifications, ancestral estimates, and the phylogeny were obtained from Kontopoulos et al. (2025). The ancestral state estimates used in the present study are based on a much larger dataset with many species lacking genomes so far, which makes such ancestral estimates more reliable than what would be obtained based on only the 190 species subset. Note that we included torpor estimates for internal nodes only if their posterior probability was above 0.95 according to a Bayesian ancestral state inference (Kontopoulos et al. 2025). This figure was rendered using the phytools R package (v. 2.4-4; Revell 2024).

These three codings represent different approaches to modelling torpor evolution and the influence individual protein-coding genes may have on it.

### Linking evolutionary shifts in torpor to gene losses and positive selection

To detect associations between individual protein-coding genes and evolutionary shifts in torpor, we first used TOGA (Kirilenko et al. 2023) to identify 19,289 one-to-one orthologs between the human genome and up to 189 genome assemblies of placental mammal species (see Methods). Human was chosen as a reference species because a) its genome is thoroughly annotated and b) humans are unable to enter torpor (the likely ancestral condition for placental mammals; Kontopoulos et al. 2025). For each orthologous gene, we identified species that likely lost the gene based on the presence of inactivating mutations (e.g., early frameshifts, stop codons), and inferred the branch(es) in the phylogeny where the loss event(s) occurred (see Methods). We also used an unbiased approach to identify phylogenetic branches with significant evidence for positive selection (Smith et al. 2015), which can indicate a change in protein function. Next, we fitted a statistical model that links shifts in torpor state with losses and/or positive selection patterns in a given gene, controlling for phylogenetic relationships. More specifically, this model treats torpor as a threshold trait, governed by an unobserved continuous variable (the “liability”; Felsenstein 2005; Hadfield 2015; see Methods). It then identifies the fraction of torpor variance explained by a particular gene, that is, the extent to which this gene contributes positively or negatively to the evolution of torpor. If a gene promotes torpor, it may be positively selected in torpor-capable lineages but lost or not under positive selection in non-torpor lineages. In contrast, genes that are generally lost in torpor-capable lineages may either inhibit torpor (in which case the gene loss is adaptive; Morris et al. 2012; Albalat and Cañestro 2016) or are obsolete once torpor has evolved (the ‘use it or lose it’ principle). In the most extreme scenario, where a gene shows a common pattern (either loss or positive selection) in all torpor-capable lineages and the opposite pattern in all non-torpor lineages, the fraction of torpor variance explained by this gene would be 1. To avoid applying the model to “invariant” genes with no or insufficient gene loss or selection signatures, we filtered our dataset down to a subset of 8,595 genes (see Methods).

After running this method for 8,595 orthologous genes and 190 placental mammals, we did not detect any protein-coding gene whose loss and positive selection patterns were strongly linked to evolutionary shifts in torpor in any of our three torpor codings. More precisely, the fraction of variance explained by individual genes was consistently below 0.2 (Fig. 2a-c, Tables S5-S7). When we examined the genes that explained the highest amount of variance for each torpor coding (Fig. 2d-f), we found that they show loss/selection signatures linked to torpor in only a few lineages (e.g., hibernating bats) and that their evolutionary patterns were not repeated in other lineages where torpor shifts also occurred. We also noticed that the explanatory power of individual genes varied considerably across our three codings of torpor. Specifically, the two torpor codings with the highest degree of similarity in genes’ explanatory power were the torpor continuum and binary torpor, with a Kendall’s τ correlation coefficient of 0.64 (Fig. S1), indicating a major effect of how torpor is coded on the results.

**Fig. 2.**
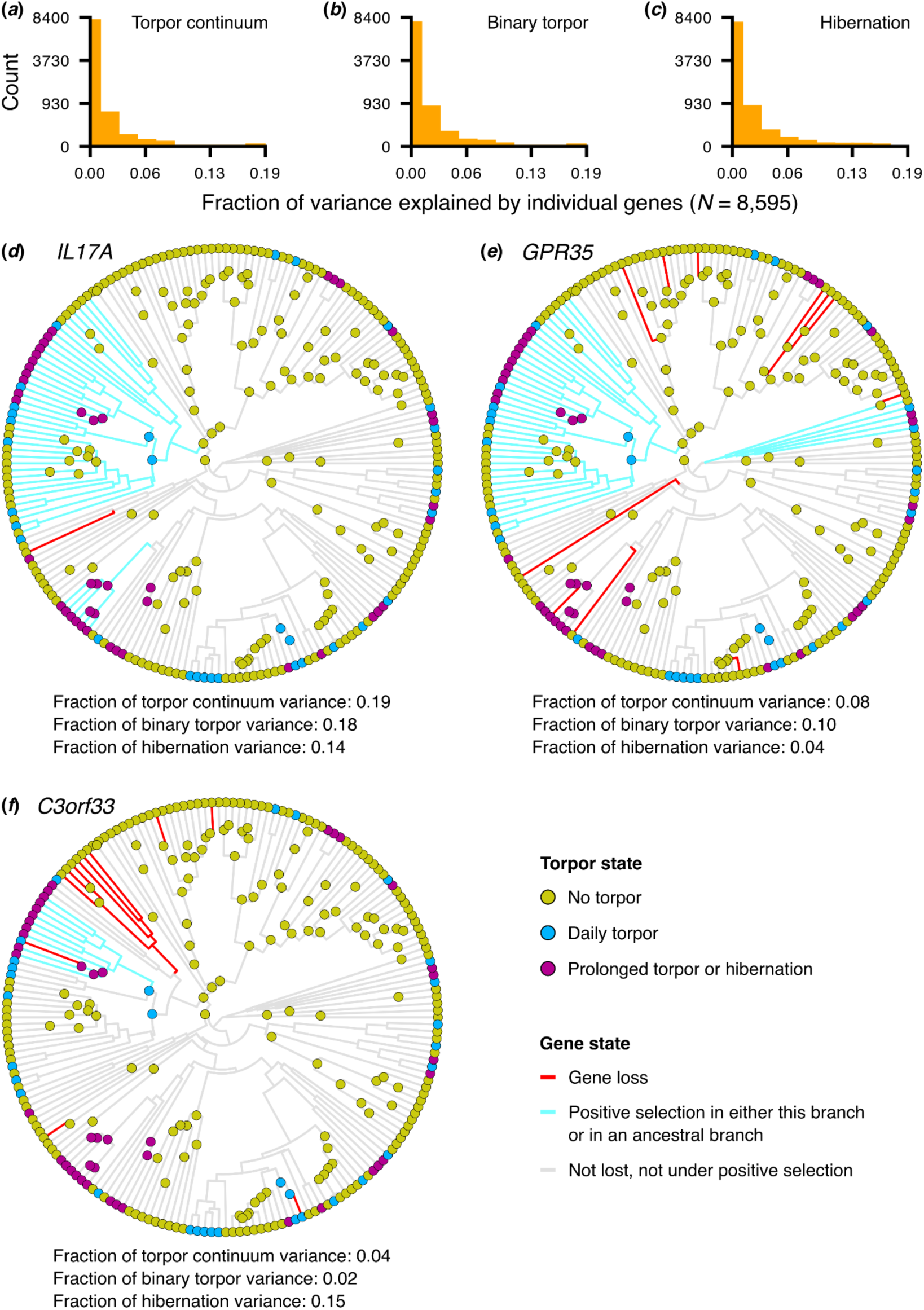
Associations between evolutionary shifts in torpor use and individual protein-coding genes, estimated through genomic screens for gene losses and positive selection across 190 placental mammals. (a-c) Histograms of the variance explained by each of 8,595 genes (after filtering for sufficient signatures of gene loss and positive selection; see Methods), separately for each torpor coding. Note that values along the vertical axes do not increase linearly. (d-f) Three representative genes that are among the top-ranked ones in at least one torpor coding; *IL17A* encoding a T-cell produced proinflammatory cytokine (d), *GPR35* encoding a G protein-coupled receptor that is expressed in the gastrointestinal tract and in immune cells (e), and *C3orf33* a less-well studied secreted protein that inhibits the AP1 transcription factor (Hao et al. 2011). *IL17A* explains the highest amount of variance (across all genes) in the torpor continuum and binary torpor screens, *GPR35* is the second most explanatory gene in the binary torpor screen, and *C3orf33* explains the highest amount of variance in the hibernation screen. Branches are coloured based on the state of each given gene (i.e., lost, under positive selection, or neither). Panels d-f were rendered using the phytools R package (v. 2.4-4; Revell 2024).

We also explored the possibility that the lack of a strong signal may be due to noise in our torpor classifications and/or genomic data. For example, certain species that are currently considered as incapable of torpor have not been thoroughly studied in conditions in which torpor would be beneficial. Similarly, torpor use by particular daily heterotherms or hibernators may have not been adequately established. To this end, we filtered our 190-species dataset down to a subset of 100 species by removing a) the lowest-quality assemblies in our list and b) species whose torpor use patterns were not well established to reduce phenotypic uncertainty (Fig. S2). For the latter, we took a very conservative approach, based on previous studies that describe species’ torpor characteristics in detail (Ruf and Geiser 2015; Nowack et al. 2023). Our filtered dataset includes 69 torpor-incapable species, 9 daily heterotherms, and 22 hibernators. Even across this carefully curated dataset, we could not identify any protein-coding gene whose loss / positive selection patterns are tightly linked to the evolution of torpor (Figs. S3 and S4, Tables S8-S10).

### Linking evolutionary shifts in torpor to changes in the rate of molecular evolution

Next, we used two alternative methods, RERconverge (Kowalczyk et al. 2019; Redlich et al. 2024) and TRACCER (Treaster et al. 2021), to investigate whether evolutionary shifts in torpor are linked to protein-coding genes exhibiting convergent changes in the evolutionary rate of torpor-capable species. We applied these methods to our curated subset of 100 placental mammal assemblies. Given that TRACCER requires at least one outgroup, we also incorporated 16 assemblies of monotremes and marsupials to the TRACCER screens (Fig. S5). TRACCER was only applied to the binary torpor and to the hibernation coding, because this method does not support traits with three (or more) ordered states.

In the torpor continuum screen, RERconverge detected only one significant gene, *ADGRF3*. This gene encodes an adhesion G-protein-coupled receptor whose function remains unclear (Lee et al. 2025), but it has previously been found to be lost in certain aquatic mammals (Sharma et al. 2018) that are incapable of torpor (Kontopoulos et al. 2025). In line with this, our screen indicates that *ADGRF3* undergoes constrained evolution during evolutionary shifts from no torpor to daily torpor (and vice versa), with its evolutionary rate not changing systematically during evolutionary shifts from daily torpor to hibernation (Fig. 3a). In the binary torpor screen, RERconverge identified *FOXP1* as the only significant gene, whereas TRACCER detected a systematic association with *UBE2W*. *FOXP1* is a transcriptional repressor that is involved in several processes, including in the development of the brain, lungs, heart, and immune system (Lozano et al. 2021; Stewart and Cho 2025). Mutations in *FOXP1* in humans can result in the *FOXP1* syndrome that is characterized by intellectual disability, developmental delays, and autism-like features, among others (Lozano et al. 2021; Stewart and Cho 2025). According to our screen, *FOXP1* exhibits higher evolutionary rates in torpor-capable lineages than in torpor-incapable ones (Fig. 3b). *UBE2W* ubiquitinates the N-terminus of substrate proteins, marking them for degradation (Scaglione et al. 2013) and generally appeared to be under constrained evolution in torpor-capable lineages and under accelerated evolution in other lineages (Fig. 4). Neither RERconverge nor TRACCER identified any significant genes in the hibernation screen. For all three genes, there are several exceptions to the patterns of sequence evolution described above. For example, there are multiple torpor-incapable lineages whose relative evolutionary rates of *UBE2W* are lower than those of certain torpor-capable lineages.

**Fig. 3.**
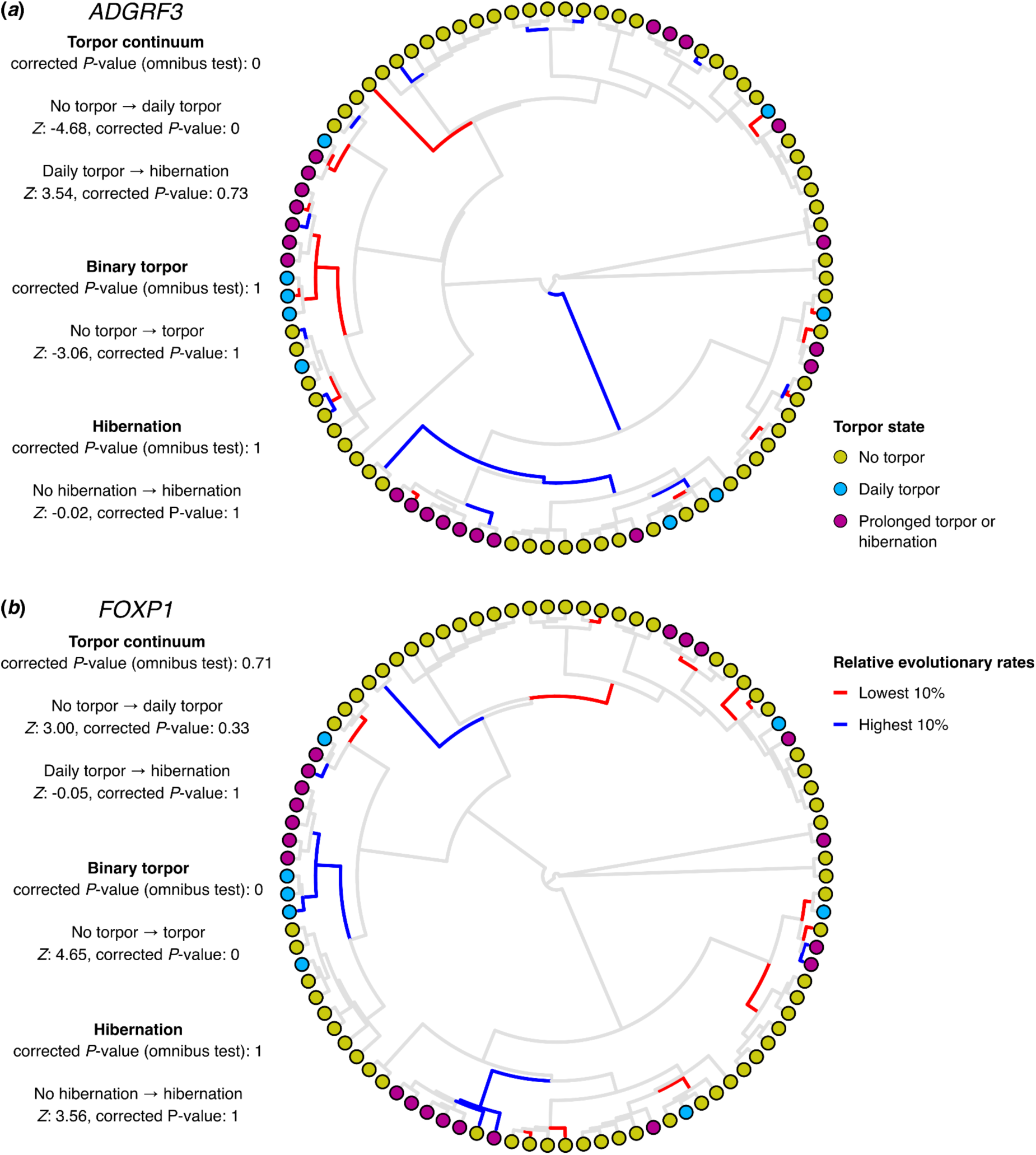
The only two genes whose relative evolutionary rates are systematically associated with our torpor codings, according to the RERconverge screen. RERconverge first examines the overall contribution of each gene to all possible evolutionary transitions among torpor states (the “omnibus” test) and then analyses each evolutionary transition separately. Positive/negative *Z* values indicate acceleration/deceleration of evolutionary rates, respectively. Note that corrected *P*-values equal to zero indicate that none of the 30,000 permulations (see Methods) could replicate (or exceed) the strength of the observed gene-torpor association. The distance of each node from the centre represents the number of tips that descend from each given node. Red and blue branches are those with the lowest 10% and highest 10% relative evolutionary rates, respectively. This figure was rendered using the phytools R package (v. 2.4-4; Revell 2024).

**Fig. 4.**
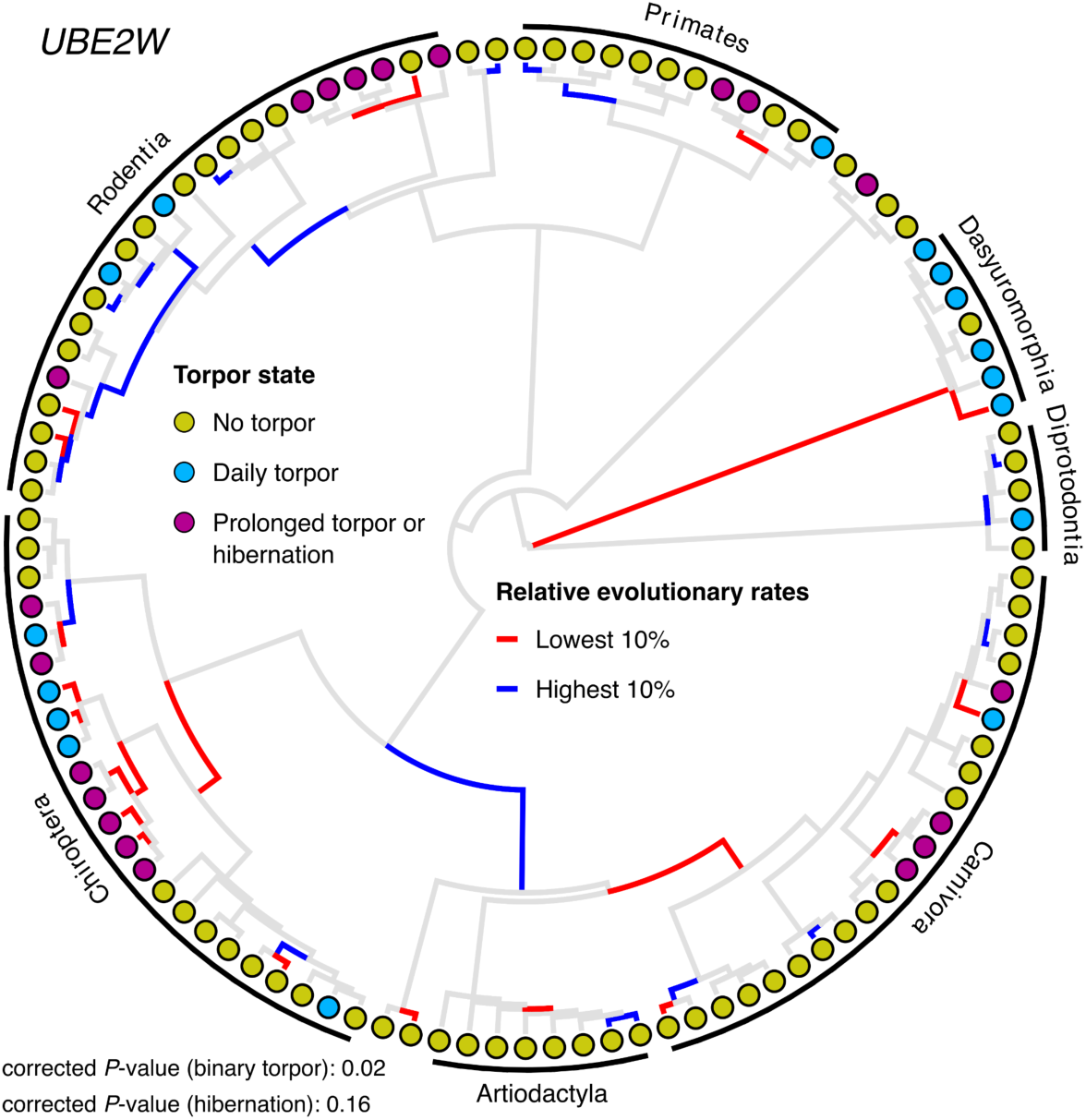
*UBE2W* is the only gene whose relative evolutionary rates are systematically associated with our binary torpor coding, according to the TRACCER screen. The distance of each node from the centre represents the number of tips that descend from each given node. Red and blue branches are those with the lowest 10% and highest 10% relative evolutionary rates, respectively. Note that all species shown in this tree (96 placental mammals and 12 marsupials treated as an outgroup) had the same transcript of the *UBE2W* gene (see Table S2). This figure was rendered using the phytools R package (v. 2.4-4; Revell 2024).

We next compared the evolutionary rate patterns in these three genes with the results from our 100-species gene loss / positive selection screen. Although *ADGRF3* shows evidence of constraint in lineages transitioning from no torpor to daily torpor, it is lost (among others) in the lineages leading to *Glossophaga soricina* (a daily heterotherm bat whose most recent ancestor in our phylogeny appears to have been incapable of torpor) and *Glis glis* (a hibernating dormouse). Positive selection in *ADGRF3* was not detected in any lineage. *FOXP1*, a gene whose rate accelerates in torpor-capable lineages, was neither lost (consistent with its functional significance) nor under positive selection in any lineage. Finally, *UBE2W* was also neither lost nor under positive selection in any lineage, despite a pattern of decreased evolutionary rates in torpor-capable lineages. Therefore, these results indicate that evolutionary shifts in torpor are not strongly tied to the evolutionary rates of individual genes across placental mammals, with any patterns of accelerated evolution likely reflecting relaxed purifying selection or drift.

### Identifying enriched pathways linked to torpor

The low explanatory power of individual protein-coding genes for evolutionary shifts in torpor raises the question of whether the evolution of torpor involved changes in distinct genes that participate in common functions and pathways. We therefore investigated whether particular pathways are systematically associated with evolutionary shifts in torpor, based on all our gene-level screens (see Methods). These analyses revealed multiple enriched pathways with corrected *P*-values below 0.05 (Fig. 5) that reflect several physiological changes that are known to occur during torpor.

**Fig. 5.**
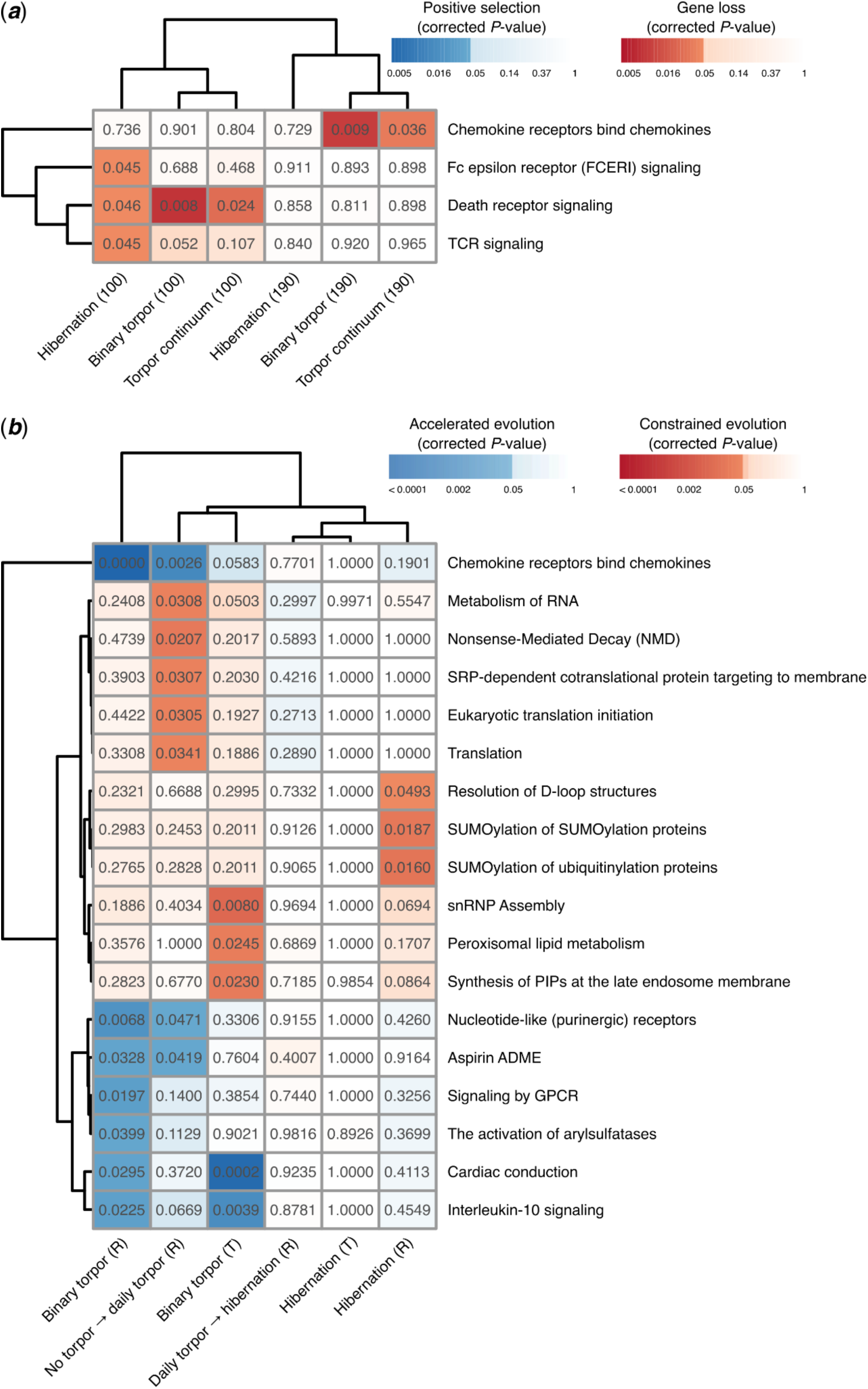
Pathways that are statistically linked to torpor evolution (rows) in one or more gene-torpor model runs (columns). Panel (a) shows pathways enriched in the gene loss / positive selection model, whereas panel (b) corresponds to evolutionary rate shift models (RERconverge and TRACCER). The numbers in each cell represent FDR-corrected *P*-values. In panel (a), the number of assemblies to which each model was applied (190 or 100) is shown in parentheses. In panel (b), “R” and “T” stand for RERconverge and TRACCER screens, respectively. Columns and rows are clustered using the complete-linkage hierarchical clustering method. Note that we applied pathway clustering to remove redundant pathways with highly overlapping gene sets (see Methods and Fig. S6 for the equivalent figure without pathway clustering). This figure was rendered using the pheatmap R package (v. 1.0.13; Kolde 2025).

Hypothermia during torpor onset is mediated by A_1_ adenosine receptor (A_1_AR) signalling in Syrian hamsters and arctic ground squirrels (Tamura et al. 2005; Jinka et al. 2011; Olson et al. 2013). Thus, our finding of acceleration in the evolutionary rate of purinergic receptors (Fig. 5b)—to which A_1_AR belongs—likely reflects a pattern of positive selection in torpor-capable species. Furthermore, torpor entrance in small mammals (e.g., hibernating squirrels) leads to a general suppression of protein synthesis, with the exception of genes involved in processes such as lipid metabolism—which becomes the primary energy source during torpor—, cryoprotection, anti-apoptotic signalling—to reduce cell death due to low body temperatures and ischemia—, and anti-inflammatory signalling (Frerichs et al. 1998; van Breukelen and Martin 2001; Heldmaier et al. 2004; Williams et al. 2005; Kurtz and Carey 2007; Shao et al. 2010; Hampton et al. 2011; Rouble et al. 2013; Vermillion et al. 2015; Logan et al. 2019; Giroud et al. 2020; Mohr et al. 2020). In contrast, hibernating black bears appear to maintain substantial levels of protein synthesis in liver and skeletal muscles to avoid muscle atrophy (Fedorov et al. 2009). Gene expression suppression in hibernating squirrels is partly achieved through changes in phosphorylation of the eukaryotic translation initiation factor 2 subunit 1 (eIF2α) and other key proteins (Frerichs et al. 1998; Miyake et al. 2015; Logan et al. 2019), which is accompanied by an increase in SUMOylation levels that confers further cytoprotection (Lee et al. 2007).

Consistent with these physiological changes, we detected evolutionary rate constraints in pathways related to mRNA processing, translation, post-translational modification through SUMOylation, and peroxisomal lipid metabolism (Fig. 5b). These pathways are likely under strong purifying selection in torpor-capable lineages due to their functional importance for torpor. Conversely, we identified gene losses and/or accelerated evolutionary rates in pathways related to apoptotic signalling and immune system function (Fig. 5a, b). The patterns in most of these pathways likely reflect a relaxation of purifying selection followed by drift that can ultimately give rise to gene losses over time. The main exception is interleukin-10 (IL-10) signalling whose rate acceleration (Fig. 5b) may arise from positive selection, given that IL-10 is an anti-inflammatory cytokine that is overexpressed during hibernation in thirteen-lined ground squirrels (Kurtz and Carey 2007). Similarly, the elevated evolutionary rate of the cardiac conduction pathway (Fig. 5b) likely reflects positive selection given that the cardiac tissue has to remain functional despite remarkable differences in heart rate between regular activity and torpor. For example, active thirteen-lined ground squirrels exhibited heart rates of up to 450 beats per minute, which were reduced to only five beats per minute during torpor (Hampton et al. 2010). Finally, our finding of evolutionary rate acceleration in arylsulfatase activation and aspirin ADME pathways (Fig. 5b) may reflect a decrease in the intensity of purifying selection in genes involved in the removal of toxic metabolites. This is in line with the downregulation of genes involved in the breakdown and removal of metabolic waste products during torpor in black bears and thirteen-lined ground squirrels (Fedorov et al. 2009; D’Alessandro et al. 2017).

Overall, there was substantial variation in pathway enrichment across different screens, with most corrected *P*-values being not much lower than 0.05. These results suggest that the evolution of torpor among placental mammals does not depend on a highly conserved set of specific gene modifications, but rather on changes in different genes within the same pathways. In other words, it appears that several lineages of placental mammals independently gained or lost the ability to enter daily torpor and hibernation (Kontopoulos et al. 2025) by modifying certain functionally important pathways—which may also vary by clade—in different ways.

### Conclusions

In the present study, we investigated whether specific protein-coding genes or pathways are linked to torpor use in placental mammals. We addressed this question by performing screens for gene losses, positive selection, and evolutionary rate shifts across a large dataset of placental mammal genomes. Despite analyzing a comprehensive set of orthologous protein-coding genes, we could not identify individual genes that are strongly linked to evolutionary shifts in torpor across placental mammals (Figs. 2–4). There was relatively stronger evidence for convergence at the pathway level (Fig. 5), which suggests that distinct but functionally similar genes may be involved in the evolution of torpor in different lineages. It remains possible that the weak associations with torpor at the individual gene level may partly be driven by noise in genome/gene alignments, gene/species’ trees, and torpor classifications, despite our best efforts to reduce such noise. Nevertheless, we should note that previous studies with analogous approaches to ours successfully detected convergent genomic signals for other (potentially less complex) traits (Indrischek et al. 2022; Morales et al. 2025; Osipova et al. 2026). Overall, our findings indicate that, at the level of individual genes, different mammalian clades likely evolved torpor through distinct genetic routes. Such heterogeneity at the genomic level is possibly responsible in part for the remarkable variation in the physiological traits, ecological strategies, and torpor patterns of torpor-capable mammals (Kontopoulos et al. 2025). Therefore, our study adds to a growing body of literature that shows that torpor may be an umbrella term, encompassing similar phenotypes across taxonomic groups that may involve different molecular components (Nowack et al. 2020, 2023; Kontopoulos et al. 2025).

A limitation of our study is that we examined the extent of genomic convergence in mammalian torpor only across protein-coding genes and not across regulatory elements. Interestingly, previous studies have provided evidence for convergent regulatory evolution across hibernators (Ferris and Gregg 2019; Nakayama and Makino 2024; Ferris et al. 2025; Steinwand et al. 2025), which may indicate that torpor evolution involves shared gene-regulatory mechanisms. However, these studies considered only a limited set of species, with the number of analyzed hibernators ranging between only four and nine. In contrast, we included 34 hibernators in our 190-species dataset (Fig. 1) and 22 in the 100-species subset (Fig. S2). Thus, the generality of reported patterns at the regulatory level needs to be addressed by future studies with larger and more taxonomically diverse datasets.

In conclusion, given the sparsity of evolutionary signatures affecting individual genes at the pan-placental mammal level, our study indicates that separate examinations of major clades (e.g., rodents) and possibly also their subclades (e.g., squirrels, heteromyids, murids) are needed to reveal genes contributing to torpor evolution. This strategy would shed light on the similarities and differences in the genes that contribute to torpor and how torpor is achieved across taxonomic groups. As high-quality genomes are rapidly becoming available, such a goal appears increasingly realistic.

## Methods

### Genome-wide screens for gene losses and positive selection

As a basis for all our comparative genomics screens, we aligned each of the 189 placental mammal genomes to the human hg38 genome assembly. We henceforth refer to the human genome as the ‘reference’ and the other placental mammals as ‘query’ genomes. For genome alignment, we used LASTZ (v. 1.04.15; Harris 2007) with parameters K = 2,400, L = 3,000, Y = 9,400, H = 2,000, and the default LASTZ scoring matrix. This combination of parameters has previously been shown to make the LASTZ algorithm sufficiently sensitive for aligning orthologous exons of distant placental mammals (Sharma and Hiller 2017). We then ran axtChain (v. 1.0; Kent et al. 2003) to chain local alignments, using the default parameters, except for ‘linearGap’, which we set to ‘loose’. We also executed RepeatFiller (v. 1.0; Osipova et al. 2019) with the default parameters to incorporate local alignments of repetitive elements. Finally, to further improve the specificity and sensitivity of the alignments, we used chainCleaner (v. 1.0; Suarez et al. 2017) with the ‘doPairs’ flag, the ‘minBrokenChainScore’ parameter set to 75,000, and all other parameters left to their default values.

To annotate genes, infer orthologs, identify gene losses and generate codon alignments, we next used the resulting alignment chains and the human GENCODE V38 gene annotation as input for the TOGA pipeline (v. 1.0; Kirilenko et al. 2023). TOGA automatically identifies orthologous protein-coding genes between the reference and query genomes, their orthology relationships (one-to-one, many-to-one, etc.), and whether they are likely intact or inactivated (i.e., a ‘loss’ or ‘uncertain loss’ according to TOGA) in each query genome. To infer the branches of the phylogeny on which a given gene was lost, we identified the earliest branches whose descending genomes all shared a loss of that particular gene (a parsimony approach). For genes with multiple transcripts, we also identified the transcript that was present and intact in most of our assemblies. The above process yielded a) a dataset of lost genes per branch of the phylogeny and b) a set of 19,289 one-to-one orthologs (Table S2). Each ortholog was lost in ∼19 out of 189 query species on average.

We obtained multiple codon alignments of intact one-to-one orthologs by running MACSE (v. 2; Ranwez et al. 2018) and cleaned the resulting alignments with HmmCleaner (v. 0.180750; Di Franco et al. 2019). Finally, we ran an unbiased screen to detect signatures of positive selection, separately for each gene and branch of the phylogeny, by executing aBSREL (Smith et al. 2015)—part of the HyPhy suite (v. 2.5.32; Kosakovsky Pond et al. 2020)—on each codon alignment. aBSREL additionally requires a species phylogeny, which we obtained by pruning the phylogeny used in Kontopoulos et al. (2025). Our criterion for the presence of positive selection in a gene and a branch was a corrected *P*-value below 0.05. For the screens with the 100-species subset, we generated separate codon alignments with MACSE and applied HmmCleaner and aBSREL as described above.

### Quantifying associations between evolutionary shifts in torpor and gene loss or positive selection

To estimate the associations between torpor and individual genes, we modelled torpor according to the phylogenetic threshold model (Felsenstein 2005; Hadfield 2015). In this model, a binary or ordered categorical trait is assumed to be driven by an unobserved continuous variable, the ‘liability’. As the liability crosses particular thresholds, the state of the trait changes (e.g., from no torpor to daily torpor). Binary traits have a single threshold that is typically set to 0, whereas, for traits with more than two ordered states, additional thresholds are estimated by the model. We, thus, fitted the following equation using the MCMCglmm R package (v. 2.34; Hadfield 2010, 2015), separately for each of our three torpor codings:

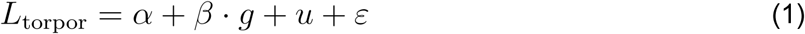

Here, *L*_torpor_ is the torpor liability, *α* the intercept, *u* a phylogenetic random effect on the intercept (based on the time-calibrated species’ tree of Kontopoulos et al. (2025)), and *ε* the residual error. *g* is a vector of the states of a given gene in each branch of the phylogeny and can take four possible values (see Fig. 2d-f and Tables S3 and S4):

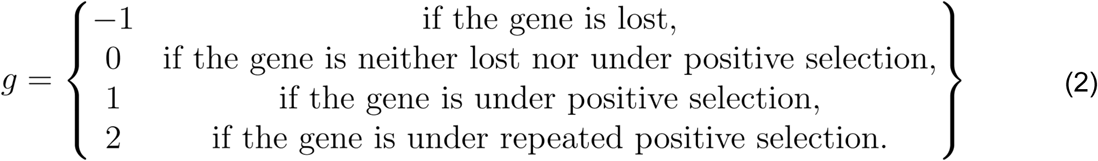

Finally, *β* is the liability-gene slope. Positive values of *β* indicate that the gene tends to be under positive selection in branches with high *L*_torpor_ values (e.g., hibernators), and vice versa.

We fitted this model separately to each gene and torpor coding. To ensure that the data are sufficiently informative to estimate model parameters, we filtered our genes dataset by excluding any genes with fewer than three unique states in *g* across all branches with a ‘known’ torpor state. This resulted in 8,595 genes being included in the 190-species screens and 5,088 genes in the 100-species screens. Regarding priors, we used the default Gaussian prior for the fixed effects and a relatively uninformative parameter-expanded prior for the phylogenetic random effect. Given that the residual variance is not identifiable in threshold models, we fixed its value to 1.

For each model run, we executed four independent Markov chain Monte Carlo chains for three million generations, obtaining samples from the posterior every 100 generations after the first 300,000 generations, which we treated as burn-in and discarded. We verified that the four chains converged on statistically equivalent and well-sampled posterior distributions by checking that, for each model parameter, the Effective Sample Size per chain was at least 1,000 and the Potential Scale Reduction Factor was lower than 1.1.

### RERconverge screens for evolutionary rate shifts

RERconverge is a versatile computational method that links evolutionary shifts in a given categorical trait to changes in the evolutionary rates of individual genes (Kowalczyk et al. 2019). For this, it calculates gene-specific relative evolutionary rates, i.e., the rates at which a gene evolves along each lineage relative to the average evolutionary rate across all genes being examined. RERconverge was recently extended to accommodate both binary and categorical traits, using continuous-time Markov models of categorical trait evolution (Redlich et al. 2024). In particular, its default approach comprises first executing a Kruskal-Wallis test (the “omnibus test”), followed by post hoc Dunn tests. The former evaluates the overall association between the gene’s relative evolutionary rate and evolutionary shifts in the trait, whereas the latter quantifies the association separately for each type of evolutionary transition between trait states (e.g., separately for transitions between a) no torpor and daily torpor and b) daily torpor and hibernation). RERconverge can also conduct phylogenetically-aware permutations (“permulations”) to correct the resulting *P*-values for the evolutionary relationships among species in the dataset. To conduct RERconverge screens, we needed i) binary or ordered categorical trait data for extant species and ii) gene trees with a common (species tree) topology.

As described above, we aligned each genome to the human genome and ran TOGA using the resulting genome alignments. Next, we generated an alignment for each gene by applying MACSE to the one-to-one orthologs detected by TOGA, including those that are likely inactivated. Non-functional orthologs should exhibit a much higher rate of evolution than their functional counterparts due to the lack of purifying selection, and, for this reason, we included them in the RERconverge screens.

To estimate gene trees, we ran RAxML-NG (v. 1.2.2; Kozlov et al. 2019), constraining each gene tree topology to that of the time-calibrated species’ tree. For each alignment, we tested four variants of the General Time Reversible DNA substitution model: a) with Γ-distributed rate variation among sites, b) with a proportion of invariant sites, c) with both of those, or d) with none of those. The optimal model per gene alignment was determined based on the Akaike Information Criterion with the small sample size correction (Sugiura 1978). To further reduce the amount of noise in the data, we excluded 4,806 gene trees in which the maximum branch length was greater than 15 substitutions per site, as such trees likely reflected highly noisy and/or gappy alignments. We then rooted each gene tree that passed the filtering based on the species’ tree.

Finally, we performed RERconverge screens for all three of our torpor codings. For torpor continuum, in particular, we provided RERconverge with a custom Markov matrix in which daily torpor lies intermediate to no torpor and hibernation, prohibiting direct transitions between no torpor and hibernation. The rate of each possible transition was estimated independently. Similarly, for binary torpor and hibernation screens, we specified the ‘all-rates-different’ model, which allows forward and reverse transitions to differ in their rate. To correct *P*-values for phylogenetic nonindependence, we conducted 30,000 permulations and applied FDR-corrections to the resulting permulation *P*-values. The number of genes analysed differed across screens, as certain genes did not have orthologs that represented all required torpor coding states. In total, 13,521 genes were included in the torpor continuum screen, 13,662 genes in the binary torpor screen, and 13,637 genes in the hibernation screen.

### TRACCER screens for evolutionary rate shifts

TRACCER is an alternative method for estimating the links between the evolutionary rates of individual genes and evolutionary shifts in a binary trait of focus. This method requires i) trait data for extant species only, ii) a species tree with branch lengths representing substitutions per site (e.g., obtained from a concatenated alignment) with one or more outgroups, and iii) gene trees fixed to the species tree topology where branch lengths represent the gene’s evolutionary rate per branch. TRACCER’s approach places greater weight on sister lineages that differ in the trait (as opposed to distant lineages) and uses a permutation strategy to assess the statistical support for the association. Compared to our gene loss / positive selection model that calculates the fraction of trait variance explained by each gene, TRACCER reports gene associations as a False Discovery Rate (FDR) corrected *P*-value.

To apply TRACCER to our curated dataset of 100 placental mammal genomes, we included 16 additional genomes of monotremes and marsupials that would form the necessary outgroups (Table S1). We obtained codon alignments and gene trees in the same manner as for RERconverge screens. We then excluded 1,862 alignments in which no monotreme or marsupial orthologs were present, and additionally 80 gene trees with maximum branch lengths of more than 15 substitutions per site

To obtain a genomic species’ tree whose branch lengths were in units of substitutions per site, we concatenated all alignments that passed our filtering criterion into a single alignment. Given that our genomes vary in their completeness and overall quality, this could potentially inflate the lengths of the branches leading to relatively lower quality genomes. To prevent this artefact, we removed all alignment columns that contained gaps using trimAl (v. 1.4.rev15; Capella-Gutiérrez et al. 2009). We then estimated a species’ tree from the resulting alignment with RAxML-NG, following the procedure described above for gene trees.

Finally, we ran TRACCER for 12 hours using 10 CPUs per screen, setting monotremes and marsupials as outgroup clades. This runtime provided sufficient permutations for each gene, with permutation counts ranging from 2,000 to more than 7 million, with a median of 4,079. TRACCER also implements a filtering strategy to exclude gene trees for which a) the number of species with or without the trait of focus is too low, b) most branch lengths are similar, or c) most branches have a length of ∼0. This ultimately led to 16,311 gene trees being evaluated in the binary torpor screen and 16,298 in the hibernation screen.

### Gene set enrichment analyses

To identify pathways that are likely linked to the evolution of torpor, we used the preranked mode of the Gene Set Enrichment Analysis software (GSEA; v. 4.3.3; Mootha et al. 2003; Subramanian et al. 2005) and the Reactome 2024.1.Hs gene set (Milacic et al. 2024). As input data, we used a) the liability-gene slopes obtained from the gene loss / positive selection model, b) the Dunn test *Z* values inferred with RERconverge, or c) the corrected *P*-values inferred with TRACCER. Because corrected *P-*values range from 0 to 1, they do not contain information—on their own—on whether a given gene is positively or negatively linked to torpor. Instead, TRACCER separately classifies each gene as ‘accelerated’ or ‘constrained’ in target lineages (e.g., in torpor-capable ones). To incorporate such information into GSEA runs, we created a new variable, *ψ*:

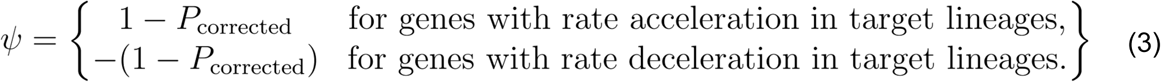

Based on Eq. (3), genes with strong statistical support for rate acceleration/deceleration have strongly positive/negative *ψ* values, respectively. For each preranked GSEA run, we used the default parameter values, except for a) the number of permutations (`-nperm`), which we set to 10,000 to increase the precision of resulting estimates and b) the gene set size thresholds, `-set_min` and `-set_max`. We set the latter two parameters to 0 and 999,999, respectively, to ensure that no gene set was excluded from the analyses.

The GSEA runs yielded pathways that were positively or negatively associated with each torpor coding and model (gene loss / positive selection, RERconverge, and TRACCER), along with key metrics such as the Normalised Enrichment Score and the FDR-corrected *P*-value. From these results, we first excluded pathways with an FDR-corrected *P*-value greater than 0.05 in all GSEA runs. To reduce redundancy, we then grouped together pathways that overlapped considerably in their gene sets. More precisely, we calculated the Jaccard similarity index for each pair of pathways and constructed a network by connecting pathways with index values of at least 0.35 (a cutoff also used by EnrichmentMap; Reimand et al. 2019). From the resulting network, we extracted connected components (i.e., pathway clusters) and, for each component, we identified the latest common parent in the Reactome hierarchy. Finally, we assigned to each parent pathway the lowest FDR-corrected *P*-value of all its child pathways.

## Acknowledgments

We thank the members of the Hiller lab for their constructive feedback on the methodology of the present study, Stephen Treaster for useful discussions regarding TRACCER, and Christoph Sinai for technical support.

## Author contributions

D.-G.K. and M.H. conceived the study and, along with A.-W.A. and B.B., designed its methodology. D.-G.K. compiled and analysed the data, generated figures, and wrote the first manuscript draft. All authors contributed to the interpretation of the results and to manuscript revisions. M.H. provided access to computational resources and supervised the study.

## Funding

Dimitrios - Georgios Kontopoulos was supported by an EMBO Postdoctoral Fellowship (ALTF 1089-2021). This work was also supported by the LOEWE-Centre for Translational Biodiversity Genomics (TBG) funded by the Hessen State Ministry of Higher Education, Research and the Arts (LOEWE/1/10/519/03/03.001(0014)/52) and a grant from the Leibniz Association’s Competition Procedure (K419/2021).

## Conflict of interest

The authors declare no conflict of interest.

## Data availability

The data that support the conclusions of this study are available as supplementary data or from Figshare at https://doi.org/10.6084/m9.figshare.30603461.

## Code availability

The code for fitting the gene loss / positive selection model and RERconverge with the custom Markov matrix is available from Codeberg at https://codeberg.org/dgkontopoulos/Kontopoulos_et_al_torpor_genomics_placental_mammals.

## Notes

https://doi.org/10.6084/m9.figshare.30603461

https://codeberg.org/dgkontopoulos/Kontopoulos_et_al_torpor_genomics_placental_mammals

### Competing Interest Statement

The authors have declared no competing interest.

### Summary of Updates

Main changes: - We have added new screens for evolutionary rate shifts linked to torpor using the RERconverge method. - We now provide a thorough discussion of how our enriched pathways are consistent with previous studies on the torpor characteristics of various species. - We reorganized the Results & Discussion and Methods sections.

