## Supplementary figures for "Lack of evidence for gene-level convergence linked to evolutionary shifts in torpor among placental mammals"

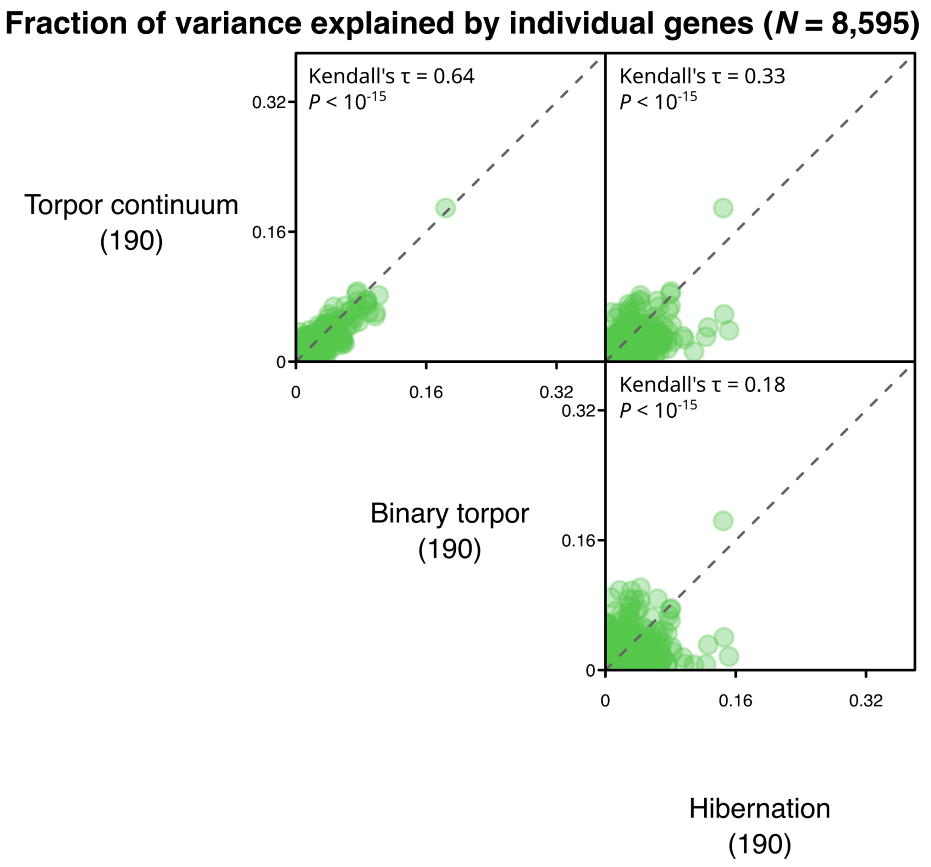


**Fig. S1.** Scatterplots of the explanatory power of individual genes across our three torpor screens, based on 190 genome assemblies of placental mammals (see figure 1 in the main text). In each panel, the dashed line stands for the one-to-one line.

**1.** Senckenberg Research Institute, Frankfurt am Main, Germany
**2.** Department of Ecology and Evolutionary Biology, University of California, Los Angeles, CA, USA
**3.** Institute of Cell Biology and Neuroscience, Faculty of Biosciences, Goethe University Frankfurt, Frankfurt am Main, Germany
**4.** School of Biology and Ecology, University of Maine, Orono, ME, USA

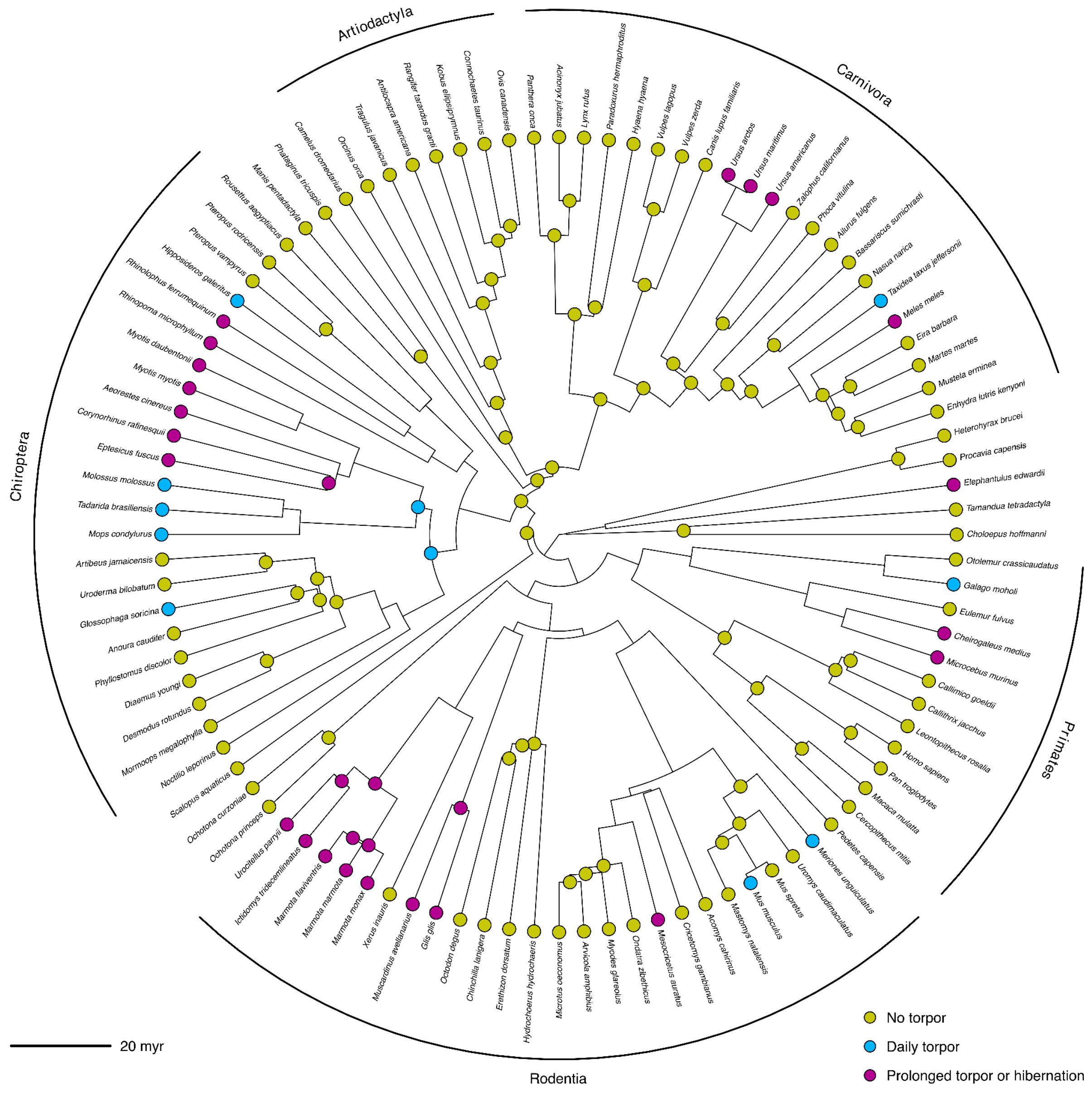


**Fig. S2.** Distribution of torpor use across the phylogeny of 100 carefully selected species of placental mammals and some of their ancestors, obtained from Kontopoulos and co-workers (see main text). The most species-rich orders in this dataset are explicitly shown. This figure was rendered using the phytools R package (v. 2.4-4).

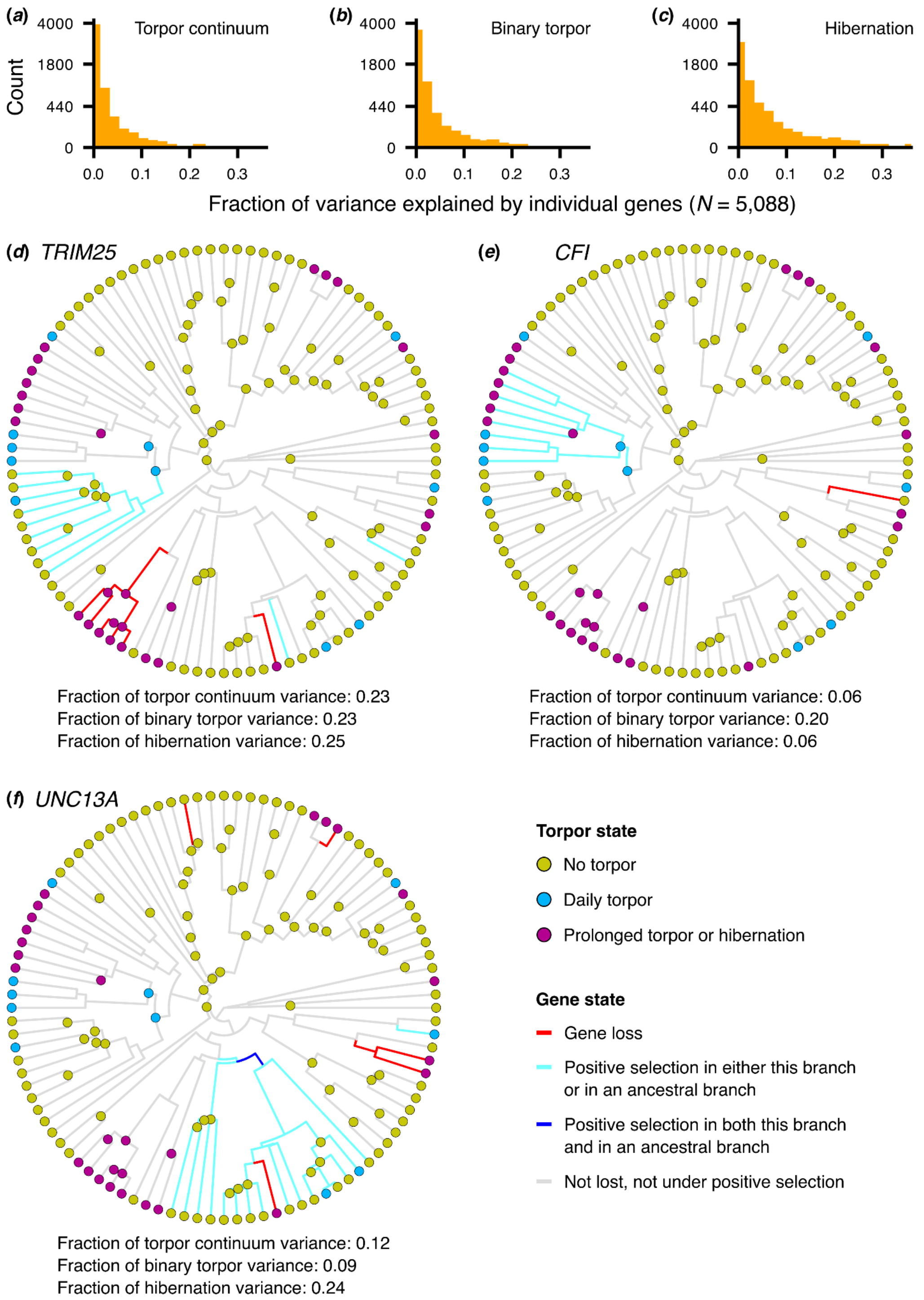


**Fig. S3.** Associations between evolutionary shifts in torpor use and individual protein-coding genes, estimated through genomic screens for positive selection and gene losses across 100 placental mammals. (a-c) Distributions of the variance explained by each gene for each torpor coding. (d-f) Three representative genes that are among the top-ranked ones in at least one torpor coding. Panels d-f were rendered using the phytools R package (v. 2.4-4). *TRIM25* explains the highest amount of variance (across all genes) in the torpor continuum, binary torpor, and hibernation screens, *CFI* is the second most explanatory gene in the binary torpor screen, and *UNC13A* is the second most explanatory gene in the hibernation screen.

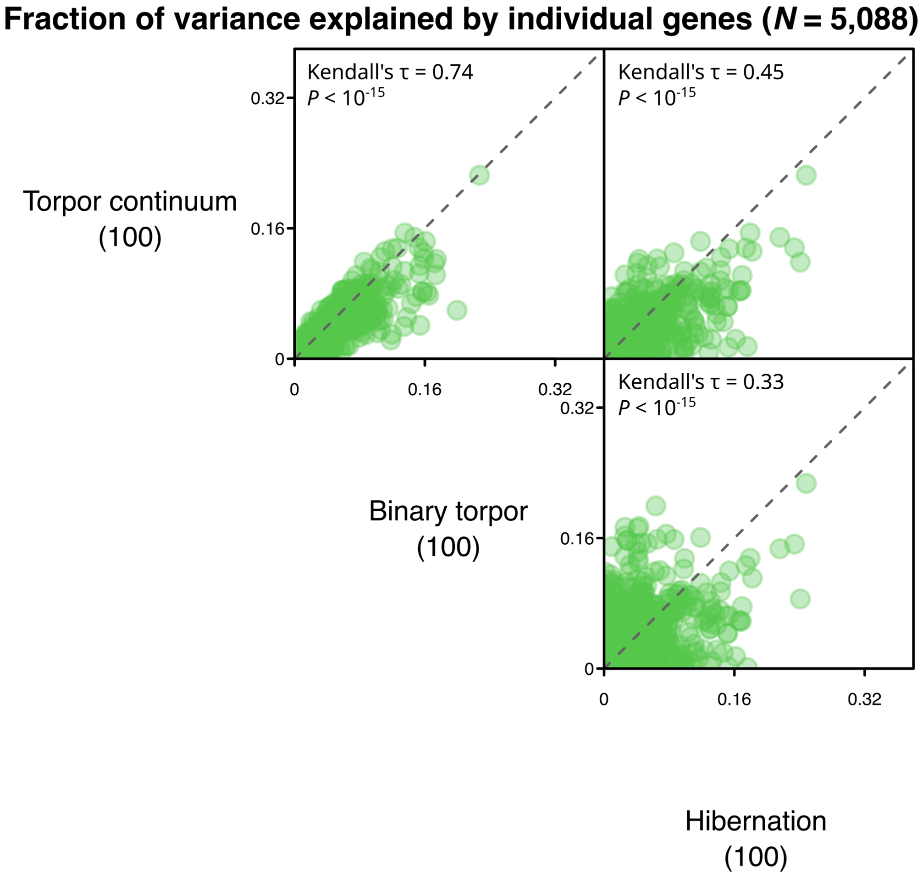


**Fig. S4.** Scatterplots of the explanatory power of individual genes across our three torpor screens, based on 100 carefully selected genome assemblies of placental mammals (see figure S2). In each panel, the dashed line stands for the one-to-one line.

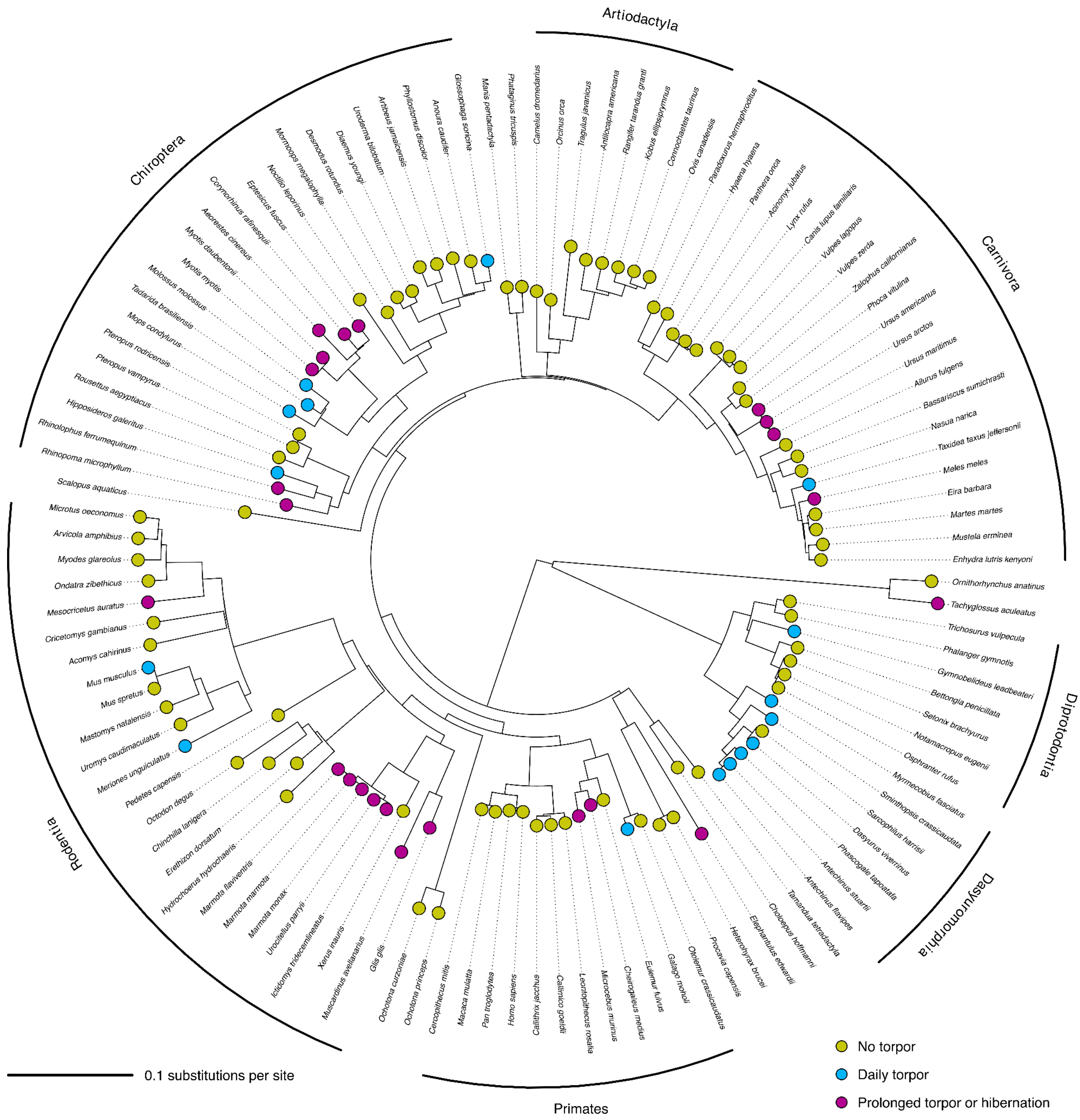


**Fig. S5.** The phylogenetic tree of the species whose genome assemblies were included in the TRACCER screens. The most species-rich orders in the tree are explicitly shown. Branch lengths were inferred with RAxML-NG, based on a concatenated sequence alignment (see Methods). The position of the root has been set here—purely for visualisation purposes—using the midpoint rooting method, as implemented in the phytools R package.

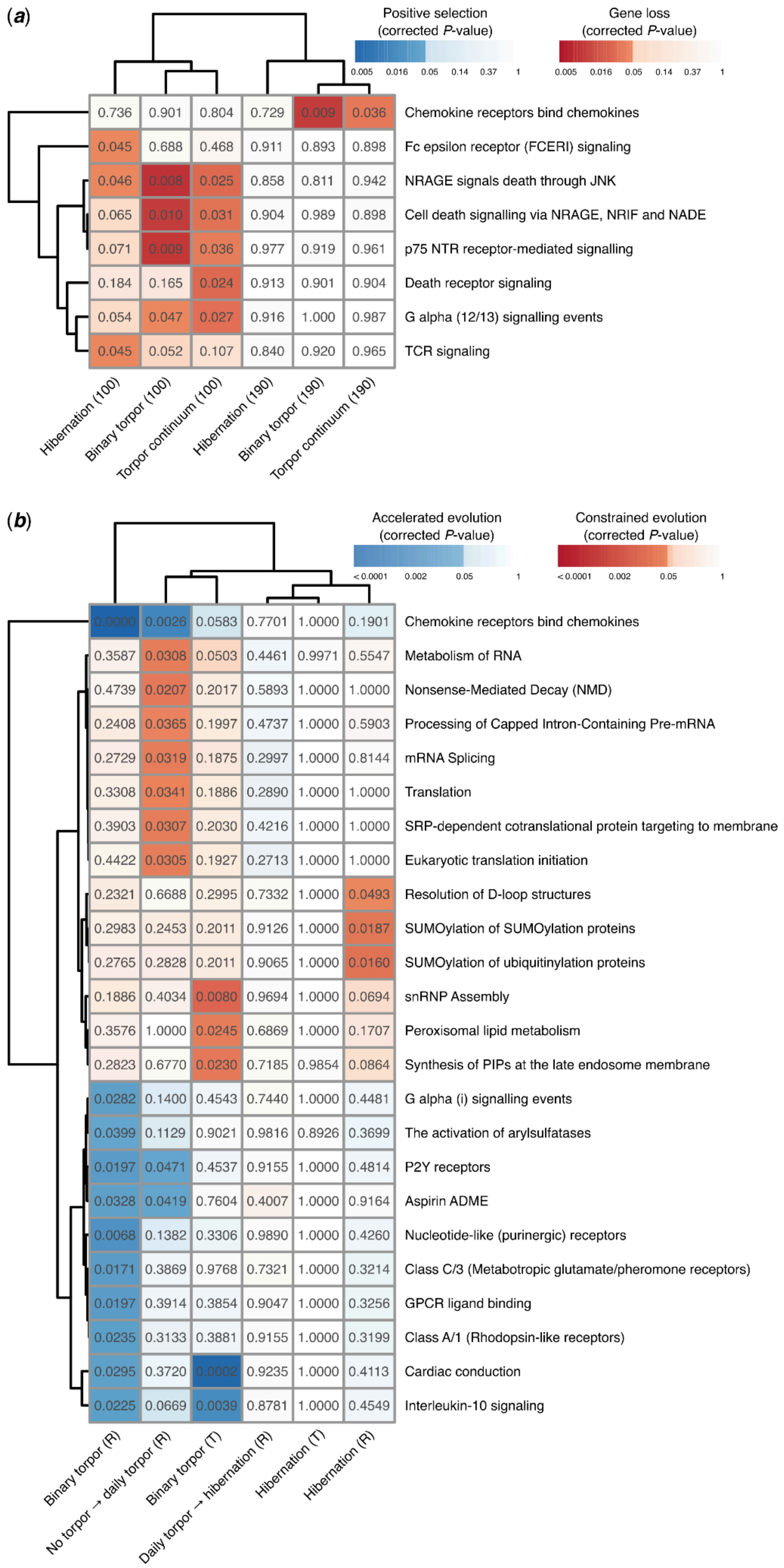


**Fig. S6.** Pathways that are statistically linked to torpor evolution (rows) in one or more gene-torpor model runs (columns). In contrast to Fig. 5 in the main text, pathway clustering has not been applied here. Thus, all enriched pathways are shown separately, including those that overlap considerably in their gene sets.
